# Executive control by fronto-parietal activity explains counterintuitive decision behavior in complex value-based decision-making

**DOI:** 10.1101/2021.11.08.467818

**Authors:** Teppei Matsui, Yoshiki Hattori, Kaho Tsumura, Ryuta Aoki, Masaki Takeda, Kiyoshi Nakahara, Koji Jimura

## Abstract

In real life, humans make decisions by taking into account multiple independent factors, such as delay and probability. Cognitive psychology suggests that cognitive control mechanisms play a key role when facing such complex task conditions. However, in value-based decision-making, it still remains unclear to what extent cognitive control mechanisms become essential when the task condition is complex. In this study, we investigated decision-making behaviors and underlying neural mechanisms using a multifactor gambling task where participants simultaneously considered probability and delay. Decision-making behavior in the multifactor task was modulated by both probability and delay. The behavioral effect of probability was stronger than delay, consistent with previous studies. Furthermore, in a subset of conditions that recruited fronto-parietal activations, reaction times were paradoxically elongated despite lower probabilistic uncertainty. Notably, such a reaction time elongation did not occur in control tasks involving single factors. Meta-analysis of brain activations suggested an association between the paradoxical increase of reaction time and strategy switching. Together, these results suggest a novel aspect of complex value-based decision-makings that is strongly influenced by fronto-parietal cognitive control.

**Highlights:**

• A value-based decision task with concurrent delay and probabilistic uncertainty
• Stronger behavioral effect of probability than delay
• Paradoxically long reaction time despite low probabilistic uncertainty
• The task activated fronto-parietal cognitive control network
• Reaction time elongation coincided with activation similar to strategy switching

## Introduction

Value-based decision-making is often difficult because participants need to evaluate a particular option not only based on its current value, but also on contextual factors that modulate its subjective value (Green et al., 2014). Even in relatively simple situations in which only a single contextual factor modulates subjective values, the participant’s decision-making behaviors often deviate from that of rational decision-makers (Chernev, 2003; Kahneman and Tversky, 1979; Simonson and Tversky, 1992). For example, when a participant is asked to choose between obtaining $100 “1 month later” or “10 years later,” they tend to select “1 month later” (Myerson and Green, 1995; Ostaszewski et al., 1998), a phenomenon known as “delay discounting” (Ainslie, 2005; Frederick et al., 2002b; Green et al., 1981; Green et al., 1994; Green et al., 1999; Kirby, 1997; Mischel et al., 1989; Rachlin et al., 1991). Similar to delay, probabilistic uncertainty to obtain the reward also affects value-based decision-making, a phenomenon known as “probability discounting” (Camerer, 1995; Green and Myerson, 2004; Kahneman and Tversky, 1979; Ostaszewski et al., 1998; Rachlin et al., 1991; Starmer, 2000; Tversky and Kahneman, 1992). For example, when a participant is asked to choose between obtaining $100 with a probability of “80%” or “10%,” they tend to choose “80%” (Ostaszewski et al., 1998). The cause of these deviations from rational decision-makers is often attributed to a tendency of human participants to minimize the required cognitive effort by controlling decision-making strategies (Basten et al., 2010; Krajbich et al., 2015; McGuire and Botvinick, 2010; Payne et al., 1963; Smith and Walker, 1993).

Value-based decision-making tasks can be even more complex in real-life settings where multiple contextual factors simultaneously modulate the values. For example, an investor evaluates the value of a company not only based on its current price but also according to uncertainty regarding its future success. At the same time, the investor needs to consider a delay to obtain a return on investment, *i.e.* the time it takes for the company to succeed, because the goal of the investor is often to maximize the profit within a given time. Recent behavioral studies have suggested that human participants can effectively handle multiple contextual factors simultaneously, but do not treat each factor equally (Vanderveldt et al., 2015). In decision-making tasks where both the uncertainty and delay of outcomes are manipulated, subjective values of given options were more strongly influenced by probability discounting than delay discounting (Blackburn and El-Deredy, 2013; Vanderveldt et al., 2015). This asymmetric processing of delay and probability suggest that cognitive processing is likely to differ between simple and complex value-based decision-making. The nature of cognitive processing and the associated brain mechanism that differentiate simple and complex value-based decision-making, however, remains elusive.

Neuroimaging studies of complex decision-making tasks have revealed that coordinated activity in the dorsolateral frontal and parietal areas enables the cognitive control necessary to handle complex task-conditions (Camilleri et al., 2018; Cocuzza et al., 2020; Dosenbach et al., 2006). On the other hand, in value-based decision-making, previous studies mostly highlighted orbital-frontal and midbrain areas related to value-coding, but not frontal-parietal areas related to cognitive control (Daw et al., 2006; Suzuki et al., 2017; Tom et al., 2007). Previous neuroimaging studies which used a complex value-based decision task (Treadway et al., 2009) also focused mostly on midbrain areas (Treadway et al., 2012; Huang et al., 2016). There are two possibilities to explain this absence of involvement of frontal-parietal areas in previous value-based decision-making studies: One possibility is that the lack of frontal-parietal activation is due to the fact that previous studies used simple value-based decision tasks in which only a single factor, such as the delay or probability, modulated the values (Hare et al., 2008; Kable and Glimcher, 2007; Tanaka et al., 2004). It is probable that frontal-parietal areas are additionally recruited in more complex value-based decision-making tasks where multiple contextual factors simultaneously affect the value. Another possibility is that value-coding in the frontal cortex and midbrain areas, by themselves, has sufficient computational capacity to calculate option values (Jimura et al., 2018; McClure et al., 2007; McClure et al., 2004). In such a case, cognitive control by frontal-parietal areas may not be necessary even in complex task-conditions in which multiple contextual factors modulate the values. In the present study, we hypothesized that the former possibility was true and devised a multifactor, value-based decision-making task which was sufficiently complex to test the hypothesis.

In the present study, we investigated the decision-making behaviors and neural activity underlying decision-making in complex conditions with multiple factors modulating the values. Our *a priori* hypothesis was that complex, but not simple, value-based decision-making recruits cognitive control mechanisms necessary to handle complex task conditions. To isolate the effect of simple and complex task-conditions, we designed a gambling task in which probabilistic uncertainty and delay to outcome was simultaneously varied (DP-task) and control tasks in which probabilistic uncertainty or delay to outcome was varied alone (D-task, P-task) (Fig. 1). The quantitative conditions presented in the D-task and P-task were matched with subsets of conditions in DP-task, which allowed us to isolate the effect of presenting multiple factors simultaneously. The brain activity in the tasks was recorded by functional magnetic resonance imaging (fMRI) to investigate the cognitive process specifically used in multi-factor decision-making.

**Figure 1.**
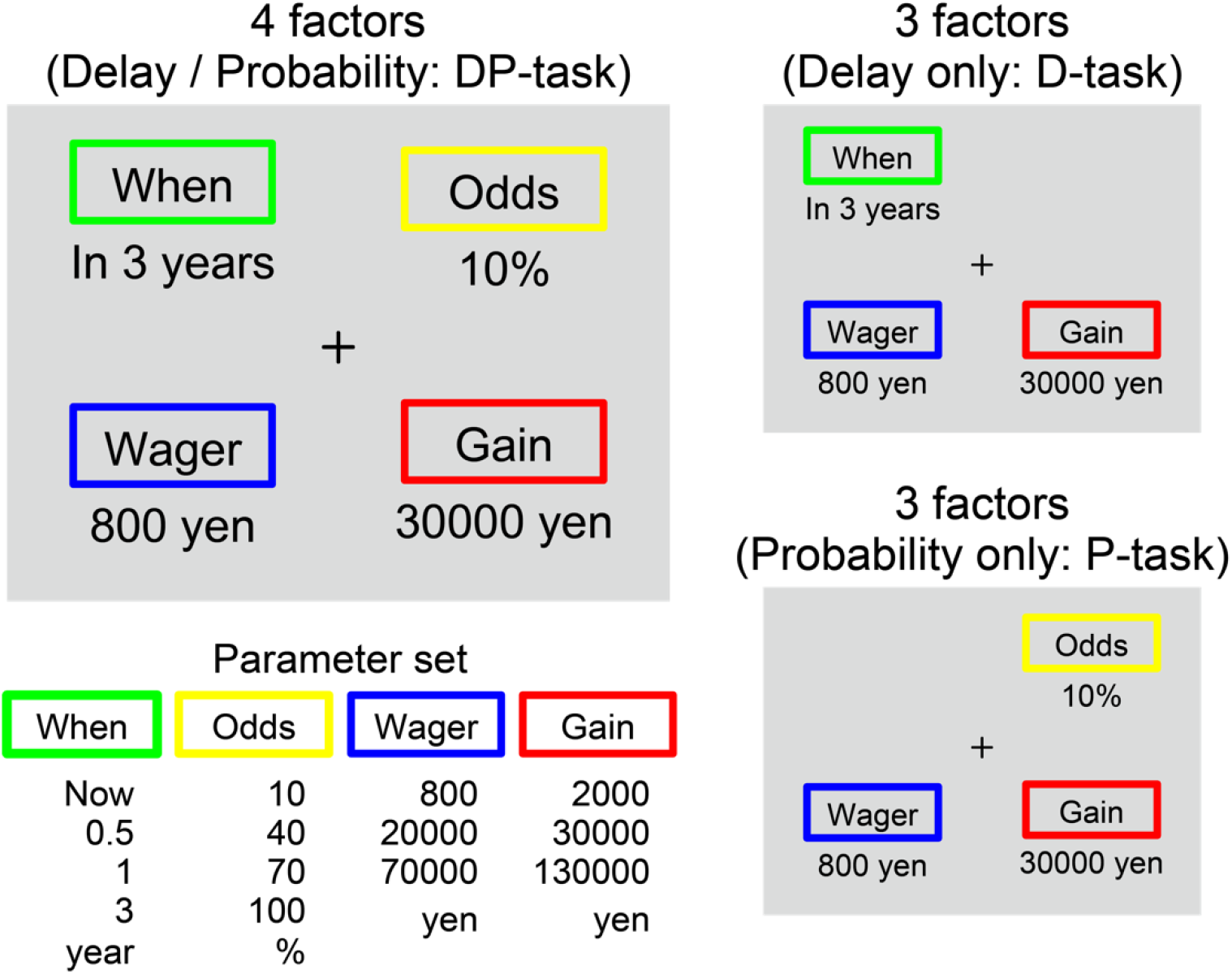
Participants performed a delayed gambling task in which they made a decision regarding whether they would accept the presented gambling condition. In DP-task (*left top*), the gamble offer involved probabilistic uncertainty (Odds), outcome delay (When), bet amount (Wager), and winning amount (Gain). In control tasks, the gamble involved only outcome delay (D-task; *right top*), or probabilistic uncertainty (P-task; *right bottom*). Parameters affecting the gamble (*left bottom*) were identical in DP-task and control tasks.

## Materials and Methods

### Objectives and hypotheses

Based on previous behavioral and neuroimaging studies, we hypothesized that cognitive processing differs between simple and complex value-based decision-making. More specifically, we hypothesized that the cognitive processing related to cognitive mechanisms is recruited in complex, but not simple, value-based decision-making. The present behavioral and neuroimaging experiments were conducted to test this *a priori* hypothesis. Based on the behavioral results, we further hypothesized, *a posteriori*, that the particular cognitive control mechanism employed in complex value-based decision-making is related to strategy switching. This latter hypothesis was examined with a meta-analysis and a functional connectivity analysis, and provided a unified interpretation of the behavioral and neuroimaging results.

### Participants

Written informed consent was obtained from 25 healthy right-handed participants (9 females; mean age, 19.5; age range, 18–22). Experimental procedures were approved by the institutional review board of Keio University and Kochi University of Technology. Participants received 2000 yen for participating.

### Behavioral procedures

Participants performed a decision-making task while fMRI was performed. In each trial, a gambling situation was presented on the screen, and participants made a judgement about whether they would accept or reject the gamble and pressed a corresponding button to indicate their decision. In the main task (DP-task), the gambling situation was defined by following factors (Fig 1 *left top*): 1) outcome delay (when the outcome of the gamble would be provided: When), 2) probabilistic uncertainty (how high chance to win the gamble is: Odd), 3) amount of bet to gamble (Wager), and 4) amount of gain when winning the gamble (Gain). Positions of the four display materials were randomized across trials. We denoted this delayed gambling task as DP-task, since the task required participants to take into account both delay and probability to make a decision. Two control tasks were also used in addition to DP-task. In one control task, probabilistic uncertainty was set as a constant (to 100%) and only delay, wager, and gain were varied (D-task; Fig 1 *right top*). In another control condition, the outcome delay was set as a constant (to “Now”) and only probability, wager, and gain were varied (P-task; Fig 1 *right bottom*). In all the three tasks, variables were drawn from the same parameter set (Fig 1 *left bottom*). Note that quantitative conditions of the trials in DP-task with 100% probability were logically equivalent to trials in D-task. Likewise, quantitative conditions in trials with immediate outcome (delay = “Now”) were logically equivalent to the trials in P-task. Participants were asked to perform the task as if the gambling conditions were real.

Trials of each task were presented in a blocked manner to minimize cognitive demand due to the change in the number of factors. The block-wise presentation of the three tasks has been shown to yield consistent results compared to an alternative presentation style where the three tasks were changed on a trial-by-trial basis (Vanderveldt et al., 2015). Each scanning session involved 2 blocks of DP-task, 1 block of D-task, and 1 block of P-task. The order of the blocks was pseudorandomized such that DP-task was unrepeated. Each task block consisted of 9 decision trials (6 sec each), 2 fixation trials (3 sec each) and 1 distractor trial (6 sec), lasting 72 seconds in total. At the beginning and the end of each block, the start and end queues were displayed for 3 seconds, respectively.

Because of the complexity of choice information consisting of 4 factors, we adopted only 4 levels for each factor in order to minimize general task difficulty. Additionally, to simplify participants’ judgement in complex decision situations, participants were required to make a decision on one choice option, whereas standard intertemporal tasks have presented two choice options simultaneously (e.g., Green et al. 1999; Green and Myerson 2004; Kable and Glimcher 2007; Vanderveldt et al. 2015, Jimura et al. 2018).

The distractor trial was imposed to ensure that the participants did not make decisions randomly without evaluating the gamble factors. Specifically, participants were presented with a gamble situation where reasonable decision was clear, and expected to reject the gamble (e.g., Wagers exceeded Gain, probability was set to 0%, and delay was set to 1000 years). The gamble stimulus set on the screen disappeared when the participant made a response.

Prior to fMRI scanning, participants received an instruction session for the tasks outside of the scanner using the actual experimental stimulus displayed on a computer monitor. They were told that the number of factors presented on the screen were 4 (DP-task) or 3 (D-task and P-task). They were also notified about the distractor trials and were instructed to reject. Then they practiced the three tasks (DP-task, D-task and P-task) for one block each. The tasks were controlled using E-Prime (Psychology Software Tools, Sharpburg PA, USA).

### Imaging procedures

A 3T MRI scanner (Siemens Verio, Germany) with a 32ch head coil mounted was used for MRI imaging. Both anatomical and functional images were acquired from each participant. High-resolution anatomical images were acquired using an MP-RAGE T1-weighted sequence [repetition time (TR) = 9.7 s; echo time (TE) = 4.0 msec, flip angle (FA) = 10°, slice thickness = 1 mm; in-plane resolution = 1 × 1 mm^2^ ]. Functional images were acquired using multi-band acceleration EPI [repetition time (TR) = 800 msec; echo time (TE) = 30 msec; number of slices = 80; slice thickness = 2 mm; flip angle = 45°; in-plane resolution = 3 x 3 mm^2^; multiband factor = 8], allowing complete brain coverage at a high signal-to-noise ratio. Each functional run involved 459 volumes (6 minutes and 7 seconds). Six runs were performed for each participant (total of 2754 volumes). The first 10 volumes of each scan were discarded to account for signal equilibrium.

### Behavioral analysis

To evaluate participants’ decision and behavior, accept rate (number of accepted trials divided by the total number of trials) and mean reaction time were calculated for each task. In DP-task, accept rate and reaction times were calculated for each combination of probability (“Odds”) and delay (“When”). Accept rate and reaction times were similarly calculated for each delay and each probability for D-task and P-task, respectively. Statistical testing of the effects of the probability and delay on accept rate and reaction times were performed by repeated measures ANOVA using SPSS Statistics 23 (IBM Corporation, NY USA). Similarly, the effect of simultaneous presentation of probability and delay was estimated using ANOVA independently for DP-task trials without uncertainty (100%) versus D-task trial, and DP-task trials without delay (Now) versus P-task.

### Image preprocessing

Image preprocessing was performed using SPM12 (http://www.fil.ion.ucl.ac.uk/spm/software/spm12/). All functional images were first temporally aligned, corrected for movement using a rigid-body rotation and translation correction, and then registered to the participant’s anatomical images. The functional images were subsequently spatially normalized to a standard MNI template with normalization parameters estimated based on the anatomical scans. The images were resampled into 2-mm isotropic voxels, and spatially smoothed with a 6-mm full-width at half-maximum (FWHM) Gaussian kernel.

### Single-level analysis

A general-linear model (GLM) approach was used to estimate task events and parametrical effects of gamble parameters. For each participant, trial events were time-locked to the presentation of the gambling situation, lasting until the participants’ response by the button press. Effects of interest were acceptance (accept or reject), tasks (DP-, P-, or D-tasks), delay (DP- and D-tasks) and probability (DP- and P-tasks). In separate GLM estimations, DP-task trials without uncertainty (100%), P-task trials, DP-task trial without delay (Now), and D-task trials were separately coded to allow direct comparison of quantitatively identical trials. Trials in DP-task without uncertainty (DP-100%) were identical to trials in D-task except that reward probability (100% in both types of trials) was explicitly presented to participants in DP-task. Similarly, trials in DP-task without delay (DP-Now) were quantitatively identical to trials in P-task except for explicit presentation of reward timing in DP-task. Brain activity was also compared between DP-100% and DP-Now using a separate GLM analysis. Trial events were then convolved with canonical HRF implemented in SPM.

We contrasted parameter estimates between 1) accept and reject trials (Fig. S2C and Table S3), 2) 100% Odds trials in DP-task vs. D-task trials (Fig. 4A and Table S4), 3) Now trials in DP-task vs. P-task trials (Fig. 4B and Table S6), and 4) 100% Odds trials and other trials in DP-task (Fig 6C and Table S7).

**Figure 4.**
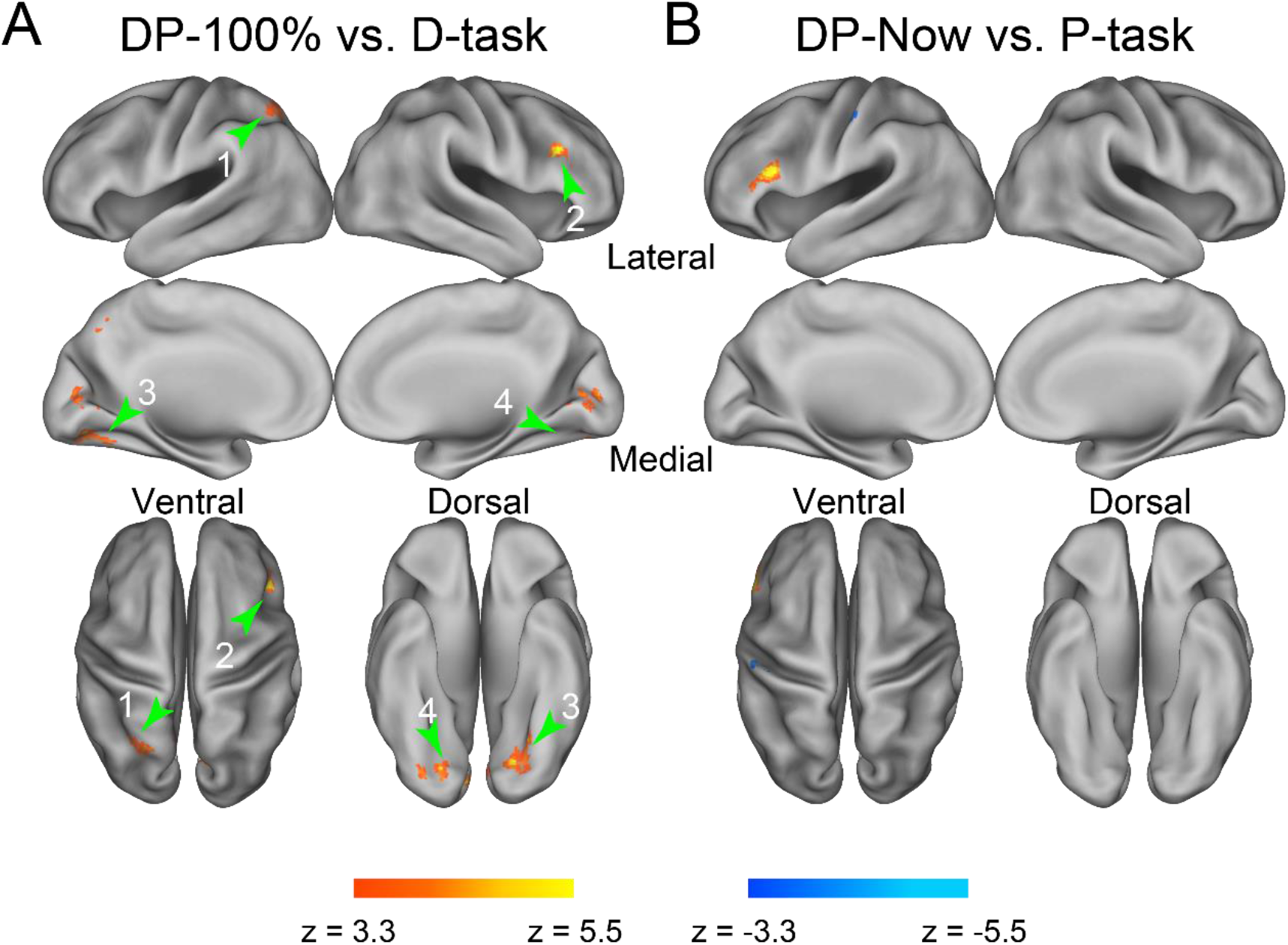
Brain activations related to processing of probability and delay in the multifactor task. A) Activation related to probability processing in multifactor context. Statistical activation maps of brain regions showing greater activity in 100% Odds trials in DP-task (warm colors) and physically equivalent control trials (D-task) (cool colors). B) Activation related to delay processing in multifactor context. Statistical activation maps of brain regions showing greater activity in Now trials in DP-task (warm colors) and physically equivalent control trials (P-task) (cool colors). 1: left superior parietal lobe; 2: right posterior prefrontal cortex; 3/4: left/right occipitotemporal cortex; 5: left posterior lateral prefrontal cortex; 6: right superior parietal cortex.

**Figure 6.**
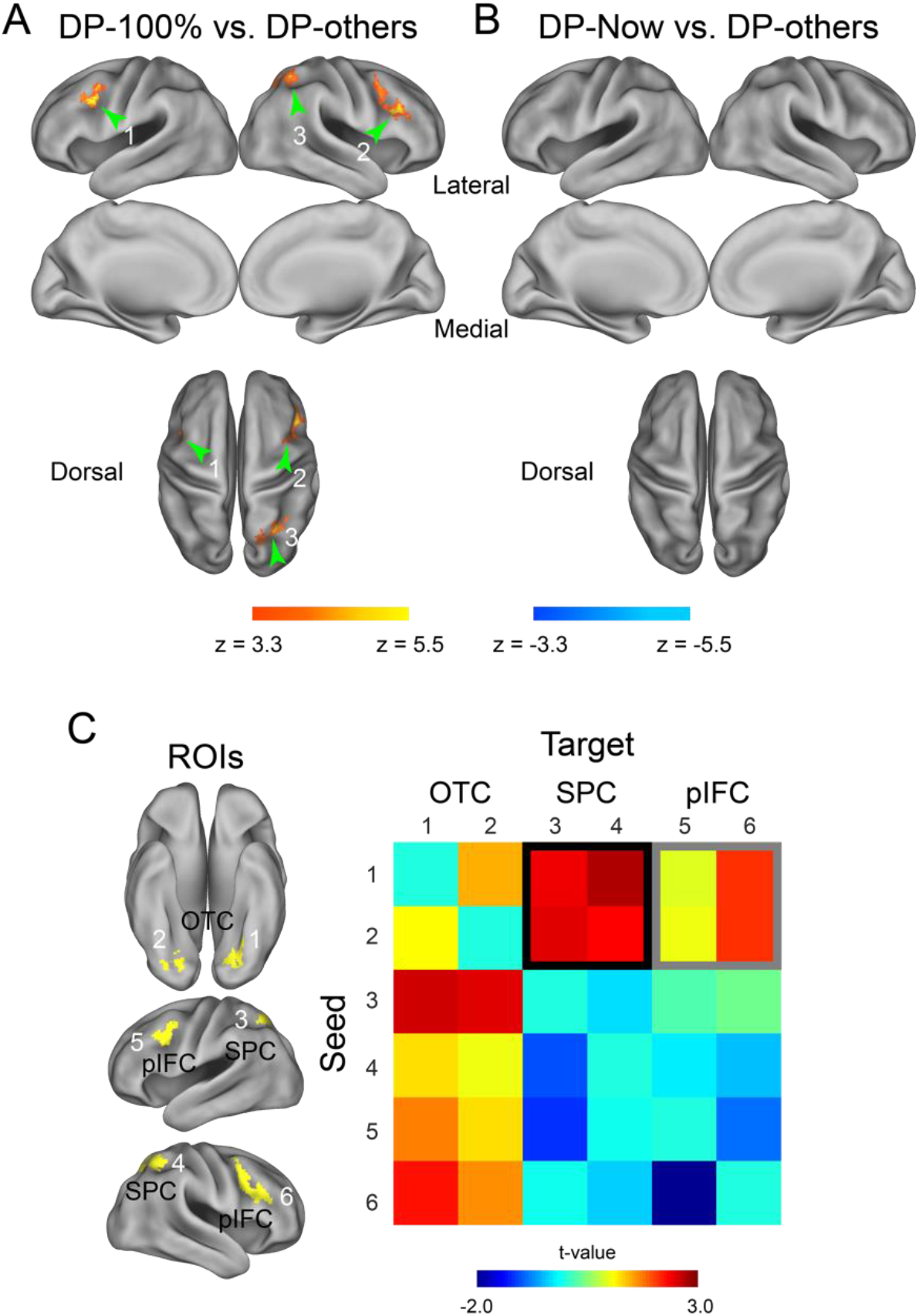
Functional interaction of brain regions related to the specific processing of 100% Odds in DP-task. A) Statistical activation maps of brain regions showing greater activity in 100% Odds (warm colors) relative to the other Odds (cool colors) in DP-task. 1/2: lateral prefrontal cortex; 3: superior parietal cortex. B) Statistical activation maps of brain regions showing greater activity in Now (hot) relative the other delays (cool) in DP-task. C) Psychophysiological interaction (PPI) analysis among activations related to the probability processing. Regions of interest were defined in occipitotemporal cortex (OTC: 1/2), superior parietal cortex (SPC: 3/4), and posterior inferior frontal cortex (pIFC: 5/6) (*left*). PPI Magnitudes in trials without probabilistic uncertainty (100%) in DP-task relative to trials in D-task were color-coded in a heat map with each cell indicating PPI between seed (raw) and target (column) ROIs. Black and gray squares indicate PPIs with P < 0.05 (corrected), and P < 0.05 (uncorrected), respectively.

For each task block, distractor trials, start cues, and end cues were also coded in GLM as nuisance regressors. Six-axis head motions, white matter, and cerebrospinal fluid (CSF) signals were also added as nuisance regressors. When extracting white matter and CSF signals, spatially normalized T1 anatomical images were segmented into white matter, cerebrospinal fluid (CSF), gray matter, bone, soft tissue, and background (air) using SPM12. Then, fMRI signal time courses were extracted using white matter and CSF images as masks.

In order to test whether the differential activity between DP-100% and D-task trials (Fig. 4A) could be explained by a general difference in cognitive load, we performed supplementary GLM analysis, where reaction times (RTs) in the DP-100% trials and D-task trials were coded as a single nuisance regressor in a GLM.

### Group-level analysis

For the group level analysis, beta maps were first contrasted within each participant and then collected from all participants. Statistical testing was performed based on nonparametric permutation testing (5000 times) implemented in *randomise* in FSL suite (Winkler et al., 2014) (https://fsl.fmrib.ox.ac.uk/fsl/fslwiki/Randomise). Clusterwise whole-brain statistical correction was performed for voxel clusters defined by a threshold (P < 0.001, uncorrected). Clusters showing significance level above P < 0.05 corrected for multiple comparisons were used as a functional mask associated the contrasts of interest described above. This group-analysis procedure was empirically validated to appropriately control false positive rates (Eklund et al., 2016). The voxel clusters listed in Tables S1-7 were also subjected to whole-brain corrections using the family-wise error rate based on the Gaussian random field theory implemented in SPM, and all clusters were significant.

### Map decoding

To characterize the current activation maps functionally, the maps for the contrasts of DP-100% vs. D-task, and trials with 100% Odds (DP-100%) vs. all the other probability conditions (10%, 40% and 70%) in DP-task, together denoted “DP-Others,” were decoded. We note that the contrast of DP-Now vs. P-task did not result in significant brain activations, hence the contrast is not described in the following.

The decoder was trained to weight a term list that characterizes a 3D brain map based on meta-analysis of functional brain mapping (https://neurosynth.org/decoder/; Yarkoni et al., 2011). High weight terms reflect greater topographical similarity between the activation maps and functional brain maps related to the words in the meta-analysis (Yarkoni et al. 2011). Full lists of weights and terms are available in Supplementary data.

Then the term lists were visualized as word clouds where the size of the words reflects the term weights. Anatomical terms and terms unrelated to brain function were excluded. Specifically, we first eliminated anatomical and general terms such as “ventrolateral,” “frontal gyrus,” and “character.” The list also included terms that are similar and/or semantically overlapping; for example, “attention” and “attentional” and “working memory” and “working.” We merged such overlapping terms into one with a weight equal to the sum of the merged terms’ weights. After this elimination procedure, we listed the top 50 terms with higher weights.

We show word clouds for the contrasts of DP-100% vs. D-task trials, and DP-100% vs. DP-others trials. Because the tested conditions (DP-100%) were identical in these clouds, we were reluctant to conduct a quantitative analysis of similarity, and instead compared the word clouds qualitatively.

### Meta-analysis maps

In order to further characterize the current activation maps, meta-analysis maps were obtained from Neurosynth (https://neurosynth.org/; Yarkoni et al., 2011). We obtained 3D maps for the search words “executive control” and “cognitive control” (P < 0.01 with whole-brain correction based on false discovery rate of uniformity test). Then, each of the two maps was binarized, and logical OR of the two binarize maps was calculated on voxel-by-voxel basis. The OR map was defined as meta-analysis mask of executive/cognitive control. This procedure was also applied to meta-analysis maps based on association tests instead of uniformity tests. The meta-analysis maps of switching with uniformity and association tests were created with similar procedure, in which “switch” and “switching” were used for the search words of Neurosynth.

### Region of interest (ROI) analysis

ROI analysis was performed to test whether the brain regions identified by the meta-analysis showed prominent activations in the current task contrasts. ROIs were defined as the meta-analysis masks as created above for executive/cognitive control and switching based on uniformity and association tests (yellow and red voxels in Figs 5B/D and S4A/4C). Then, for each ROI, signal magnitudes of the contrasts of DP-100% vs. D-task and DP-100% vs. DP-Others were calculated using *fslmeants* implemented in FSL.

**Figure 5.**
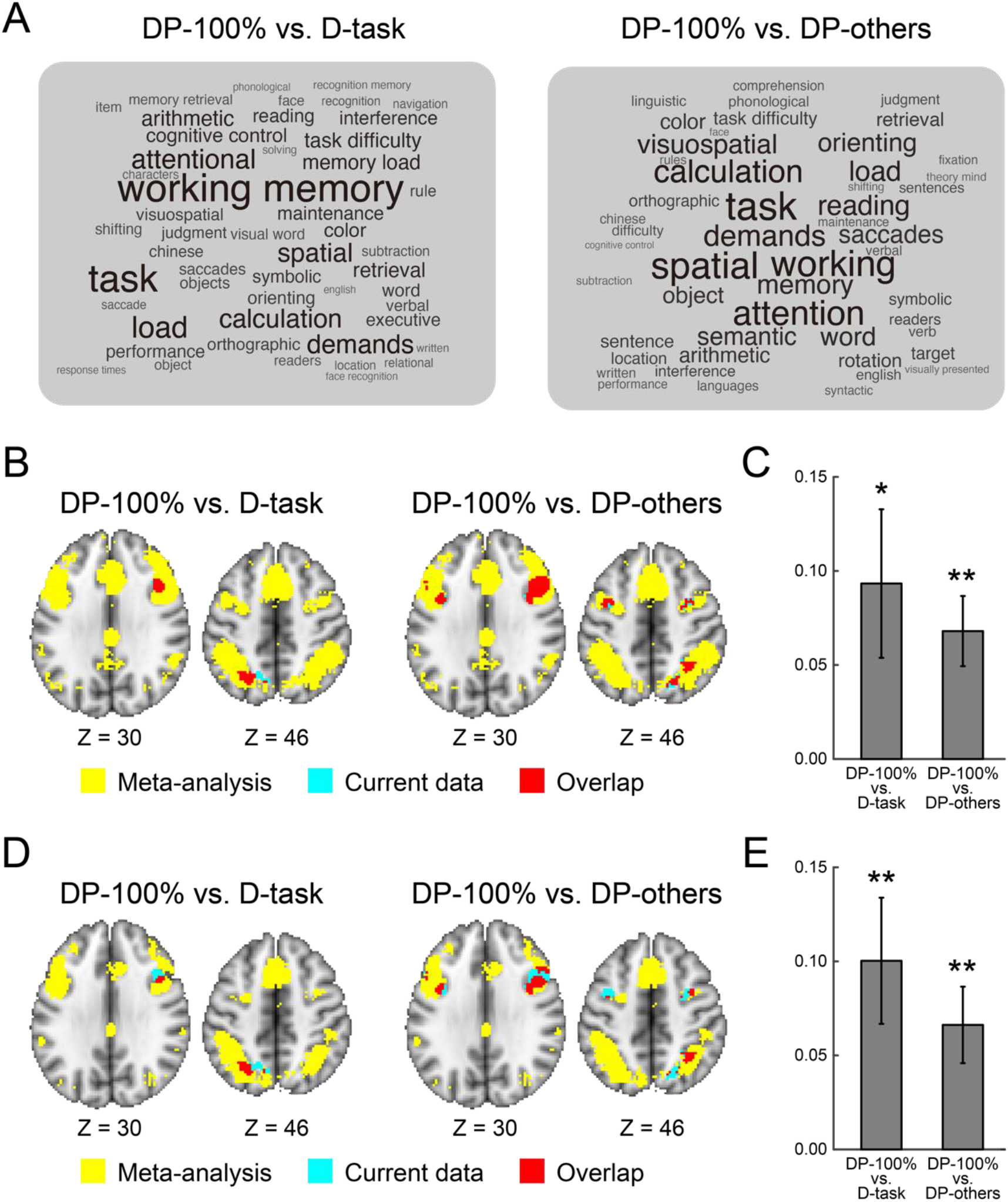
Characterization of activations by large-scale meta-analyses. A) Activation maps were decoded into functional terms, and the term lists are visualized as word clouds where the size of the words reflects the word weights. DP-100% vs. D-task (*left*), DP-100% vs. DP-Others (*right*). B) Maps of meta-analysis for cognitive/executive control are overlaid onto 2D transverse slices in yellow. The levels of the slices are indicated by the Z coordinates of the standard brain. The activation maps are further overlaid onto the slice for the contrasts DP-100% vs. D-task (*left*), and DP-100% vs. DP-Others (*right*) in cyan. The area overlapping between the meta-analysis maps and contrast maps are colored in red. C) Region of Interest (ROI) analysis. ROI was defined based on the meta-analysis maps in panel B, and signal magnitudes of the contrasts DP-100% vs. D-task and DP-100% vs. DP- Others were calculated within the ROI. Error bars indicate standard errors of the mean across participants. *: P < 0.05; **: P < 0.01. D) Maps of the meta-analysis for switching are overlaid onto 2D transverse slice in yellow. Other formats are identical to those in panel B. D) ROI analysis. ROIs were defined based on the meta-analysis maps in panel D, and other formats were identical to those in panel C.

### Psychophysiological interaction analysis

In order to examine task-related interregional interactions among brain regions during multifactor decision-making, psychophysiological interaction (PPI) analysis was performed using SPM 12 (Friston et al., 1997). The current analysis focused on brain regions and their connectivity involved in strategy switching (see Results). ROIs were defined as voxel clusters showing statistically significant activation regions in at least one of either DP-task trials without uncertainty (100%) versus other DP-task trials (Fig 6A) or DP-task trials without uncertainty (100%) versus D-task trials (Fig 4A). Then, a total of six regions of interest (ROIs) were obtained: bilateral lateral prefrontal cortex, bilateral superior parietal lobe, and bilateral occipitotemporal cortex (Fig 6C, *right*). We defined the ROIs based on the contrast that showed behavioral effect (DP-100% vs D-task).

For each ROI, the signal time course was extracted as the first eigenvariate of the voxel clusters. The percentage variances explained by the first eigenvariates were 66.29 ± 9.33 (mean ± SD) in the left Occipital Temporal Cortex (OTC) ROI, 64.26 ± 8.65 in the right OTC ROI, 75.98 ± 7.63 in the left SPC ROI, 69.85 ± 7.15 in the right SPC ROI, 66.08 ± 7.01 in the left posterior Inferior Frontal Cortex (pIFC) ROI, and 64.00 ± 9.19 in the right pIFC ROI.

A psychological variable was defined as a time series of contrast of interest, DP-task trials without uncertainty versus D-task trials. An interaction effect of the seed time course and psychological variable was calculated based on SPM12. The interaction effect, the psychological variable, and the timecourse of the seed region were included in GLM. As additional nuisance effects, nuisance behavioral events, six-axes head-motion, and time courses of white matter signal and cerebrospinal fluid signal were also included in GLM. Then, voxel-wise GLM estimations were performed for all other ROIs (target), and beta-values were averaged within each ROI for each participant. Finally, averaged beta-values of PPI were collected for all combinations of seed/target ROIs (total 30) from all participants, group-level effects were tested.

For statistical testing, PPIs between seed and target regions were calculated using *fslmeants*, and averaged across contralateral and ipsilateral hemispheres, as we did not observe strong hemispheric asymmetry in PPIs (Fig 6C; Misonou and Jimura 2021). Then, the significance of the PPI strength was tested by the one-sample t-test. P-values were corrected for multiple comparisons based on Bonferroni correction.

## Results

We collected fMRI data while participants decided whether to accept or reject gambles that offered a chance of gaining or losing money. For each trial, the chance of gaining money and the time to outcome were varied simultaneously and independently. In each trial, the price for betting (“Wager”), the amount of gain (“Gain”), the probability of winning (“Odds”), and the time to outcome (“When”) were presented on the screen (DP-task) (Fig. 1). Participants were asked to decide whether to accept the delayed gamble and then indicate their decisions by pressing a button. In two control tasks, either probability or delay to outcome was set as a constant and excluded from the offer (D-task, P-task; Fig. 1).

The accept rate for DP-task was higher for conditions with a smaller Wager [t(24) = −5.1, P < 0.001] and a greater Gain [t(24) = 2.2, P < 0.05] (Fig. S1), confirming that participants made decisions based on the presented amount of Wager and Gain. For the same Wager and Gain, participants accepted the delayed gamble more when the Odds were higher (Fig 2), consistent with probability discounting (Green and Myerson, 2004; Kahneman and Tversky, 1979; Tversky and Kahneman, 1992). Repeated measures analysis of variance (ANOVA) with 4 levels of Odds and 4 levels of When as factors revealed a significant positive effect of Odds on the accept rate [F(1,24) = 459.8; P < 0.001; with linear contrast of Odds]. Similarly, participants accepted the delayed gamble more if When was shorter [F(1,24) = 6.2; P < 0.05; with linear contrast of When; Fig. 2], consistent with delay discounting (Frederick et al., 2002a, b). The interaction between Odds and When was not significant [F(1,24) = 3.5; P = 0.08; with linear contrast of an interaction of Odds and When]. Thus, the level of statistical significance was higher for the probability than the delay, consistent with previous studies reporting that the effect of probability was stronger than that of the delay when presented together (Blackburn and El-Deredy, 2013; Vanderveldt et al., 2015). These results collectively suggest that participants performed the decision-making task in a value-based manner despite the complexity of the task structure.

**Figure 2.**
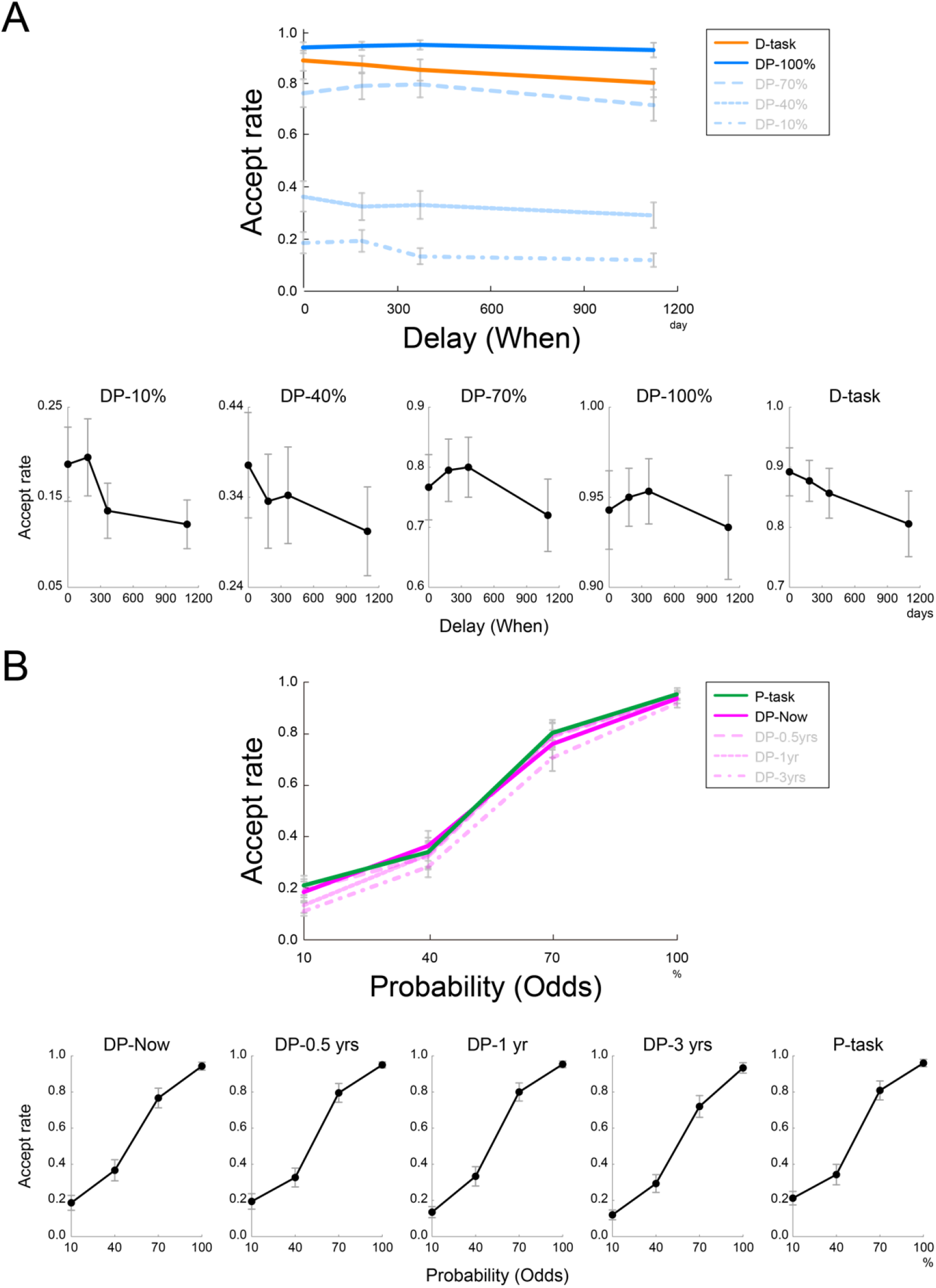
Accept rates as a function of delay (“When” parameter; A) and probability (“Odds” parameter; B). The tasks (DP-, D-, or P-task) and task parameters (delay and probability) are indicated on the right. Plots are magnified for each condition at the bottom. Error bars indicate standard errors of the mean across participants.

We next examined whether or not probability and delay affected participants’ decisions differently when both factors were presented together versus when each factor was presented alone. To isolate the effect of the simultaneous presentation, we set the quantitative conditions of the control tasks (D-task and P-task) equivalent to subsets of conditions in the DP-task: Conditions in the D-task were quantitatively equivalent to the conditions with “Odds = 100%” in the DP-task. Similarly, conditions in the P-task were quantitatively equivalent to the conditions with “When = Now” in the DP-task (unless otherwise noted, comparisons between the DP-task and the D-task or P-task were done using the quantitatively equivalent conditions). Comparing the D-task and DP-task revealed significantly larger accept rates for the latter [t(24) = 2.47; P < 0.05; two rightmost panels in Fig. 2A], suggesting that participants’ decisions were altered by explicit presentation of the probability information in addition to the delay information. In contrast, comparing the P-task and DP-task showed no significant difference in accept rate [t(24) = 1.11; P = 0.28; rightmost and leftmost panels in Fig. 2B]. These results indicated that explicit presentation of probability information (100%), but not delay information (Now), altered the participants’ decision to accept the offer, suggesting that the probability but not delay, was processed differently depending on whether the two factors were presented together in the DP-task.

To further characterize the decision behavior, we next compared the reaction times in the DP-task with those in the control tasks. When performing a cognitively demanding task, reaction time tends to increase as a function of the number of factors that the participant needs to take into account (Treisman, 1993; Woodman and Luck, 2004). Reaction time data were not skewed at any levels of Odds and Delay in the DP-, D-, and P-tasks (zs < 1.50; P > 0.14). Consistently, compared with the D-task, the reaction time was significantly longer in the DP-task [t(24) = 4.35; P < 0.0001; two rightmost panels in Fig. 3A]. Similarly, the reaction time was significantly longer in the DP-task than in the P-task [t(24) = 6.51; P < 0.001; two rightmost panels in Fig. 3B], although the accept rates were equivalent. Thus, the overall increase in reaction time occurred both for probability and delay, suggesting that an increase in the number of factors does not simply explain the increase in the accept rate specifically for probability.

**Figure 3.**
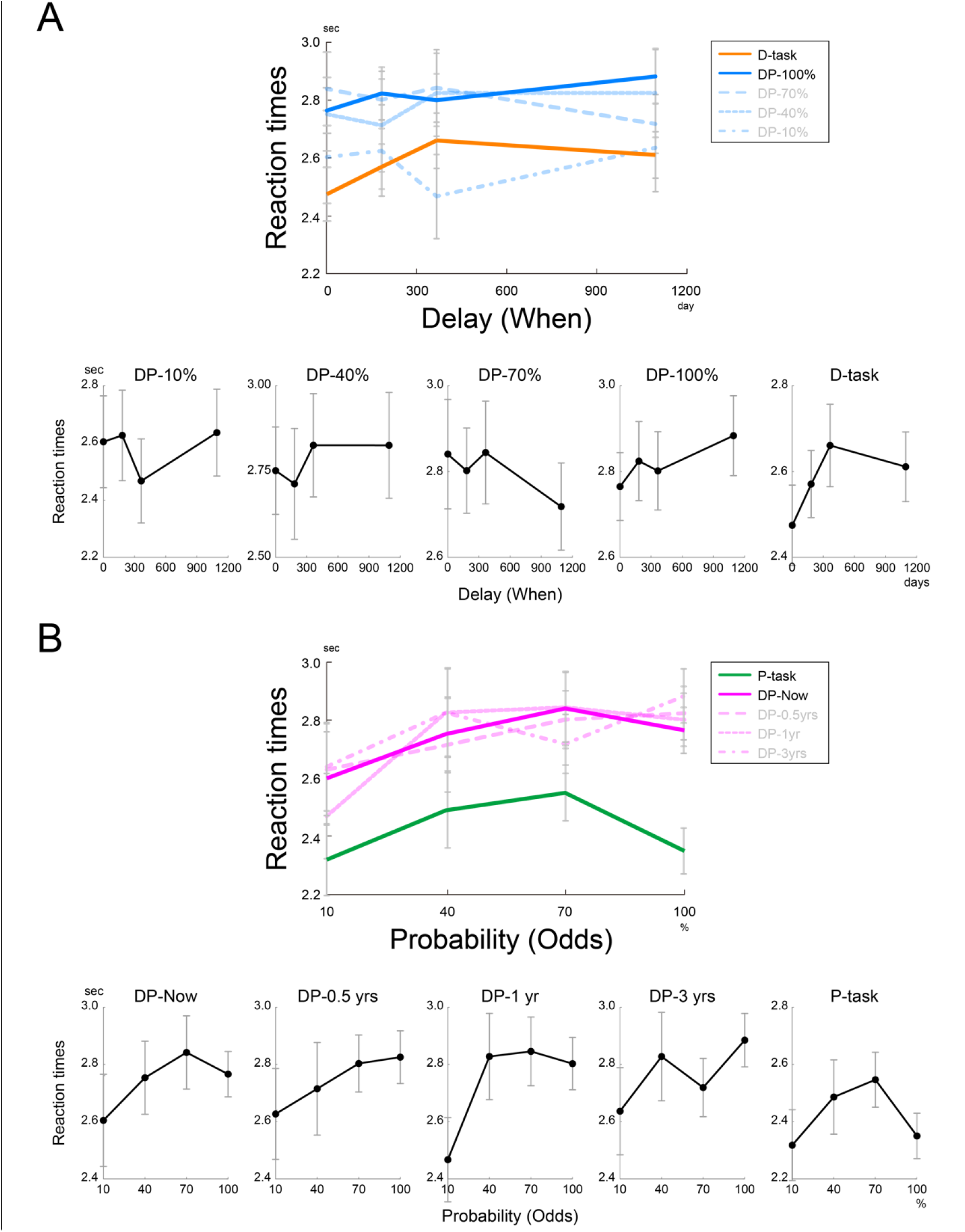
Reaction times as a function of delay (“When” parameter; A) and probability (“Odds” parameter; B). The tasks (DP-, D-, or P-task) and task parameters (delay and probability) are indicated on the right. Plots are magnified for each condition at the bottom. Error bars indicate standard errors of the mean across participants.

Despite the similar increase in overall reaction time, precise relationships between reaction time and each factor in the DP-task and control tasks revealed a difference between probability and delay. Of interest, one-way repeated measures ANOVA with 4 levels of Odds as a factor revealed that reaction time was an inverse U-shape as a function of Odds in the P-task [F(1,24) = 10.77, P < 0.01; with a planned contrast in which the middle two revels (40% and 70%) were greater than the two end levels (10% and 100%); the rightmost panel in Fig. 3B]. This was consistent with previous reports that a participant’s decision is faster when the uncertainty is lower (Bestmann et al., 2008). Indeed, compared with high uncertainty conditions (*i.e.* 40% and 70% Odds), the reaction time was significantly shorter in both the 10% Odds [t(24) = 2.56, P < 0.05] and 100% Odds conditions [t(24) = 2.23, P < 0.05].

Then, to examine the effects of Odds and When in the DP-task, a two-way repeated measures ANOVA was performed with 4 levels of Odds and 4 levels of When as factors. The main effect of Odds on reaction time was statistically significant [F(1,24) = 4.51, P < 0.05; with the planned contrast of the inverse U-shape]. However, the relationship between Odds and reaction time did not show a clear inverse U-shape (Fig. 3B). Specifically, the reaction time was similar for high uncertainty conditions and the 100% Odds condition [t(24) = 0.35, P = 0.77], whereas the reaction time in the 10% Odds condition was markedly shorter [t(24) = 3.01, P < 0.01]. In contrast to probability, the delay parameters did not show significant effect on reaction time [linear contrast: F(1, 24) = 0.50, P = .49; inverse U-shape effect: F(1, 24) = 0.19, P = .67]. Thus, these behavioral results suggest that probability, but not delay, was processed differently in the multifactor task relative to the single factor control task. Moreover, the analyses of reaction time suggested that relative to the rest of the conditions, participants underwent additional and parallel decision processing when the 100% Odds condition was presented explicitly in the DP-task.

Because the prolonged RT in the DP-100% trials by itself does not directly address change in decision processes in those trials, we examined whether decisions were made differentially depending on the probability level in the DP-task. For each Odds level, we performed a logistic regression analysis where choice was predicted by Wager, Gain, and When parameters. In the trials involving probabilistic uncertainty (i.e., 10%, 40%, and 70%), Wager showed a significant effect on acceptance [10%: t(24) = −3.64, P < 0.01; 40%: t(24) = −4.41, P < 0.001; 70%: t(24) = −2.33, P < 0.05], indicating that participants rejected gambles more frequently in trials with higher Wager. Gain also showed significant effect on acceptance [10%: t(24) = 2.40, P < 0.05; 40%: t(24) = 3.65, P < 0.01, 70%: t(24) = 2.21, P < 0.05], indicating that they accepted gambles more frequently in trials with higher Gain. On the other hand, in trials without probabilistic uncertainty (i.e., 100%), neither Wager nor Gain showed a significant effect [Wager: t(24) = −0.40, P = 0.69; Gain: t(24) = 1.22, P = 0.23]. Notably, these beta coefficients of Wager were significantly different between certain and uncertain trials [100% vs. 10%: t(24) = 3.50, P < 0.01; 100% vs. 40%: t(24) = 4.34, P < 0.001; 100% vs. 70%: t(24) = 2.32, P < 0.05]. For Gain, the beta coefficient differed between 100% and 40% probability trials [t(24) = −2.92, P < 0.01]. The differential coefficients clearly demonstrated that participants used different acceptance strategies depending on probabilistic uncertainty. Specifically, when the gamble involved probabilistic uncertainty, participants considered Wager and Gain as aversive and preferred factors, respectively, whereas in the 100% Odds trials, such consideration was absent.

Based on these behavioral results, we next proceeded to examine brain activations. Consistent with the larger behavioral effect of probability in DP-task, processing of probability recruited larger and more widespread brain activations than the processing of delay (Fig S2A-B; Tables S1/2). These parametrical effects did not differ between DP-task and control tasks, suggesting that the parametrical effects were comparable in those tasks. To further identify brain regions specifically recruited during multifactor decision-making, we examined DP-task and control tasks which had physically equivalent conditions (Fig 4). In particular, we subtracted activation maps during D-task from those during DP-task with 100% Odds (DP-100%), aiming to isolate additional processing of probability information in the multifactor context. The comparison revealed widespread brain activations in the pIFC, superior parietal lobe (SPL), and OTC (Fig 4A; Table S4). In a separate GLM analysis, where reaction times in each of DP-100% and D-task trials were coded as a separate parametric effect (see Materials and Methods), these activations were also observed (Fig S3 and Table S5), suggesting that these activations were not simply explained by the difference in general cognitive load between these trials.

In contrast, a comparison between the P-task and DP-task with immediate outcome (DP-Now), which isolated the additional processing of delay information, revealed much smaller brain activations within the inferior frontal sulcus, the more ventral anterior part of the IFC (Fig 4B and Table S6). This pattern of brain activations closely parallels the result that a difference in accept rate between the multifactor task and the control task was seen for probability but not for delay (Fig. 2A-B).

One important question about the activation map in DP-task vs. D-task (Fig. 4A) is what cognitive functions the activation map may reflect. To search for potentially relevant cognitive functions, we conducted a decoding analysis such that the activation maps were labeled as word clouds based on their topographical similarity to functional brain maps in meta-analyses (see Materials and Methods). The word clouds primarily included cognitive terms related to executive and cognitive control such as working memory, attention, and task demands (Fig 5A; see also Supplementary data for full list of words), which is implemented in fronto-parietal mechanisms (Corbetta and Shulman, 2002; D’Esposito and Postle, 2015; Dosenbach et al., 2006; Miller and Cohen, 2001). The decoding results suggest that the fronto-parietal involvements in DP-task vs. D-task (Fig. 4A) reflected executive and cognitive control functions. To test this possibility more specifically, we examined spatial characteristics of our results and the meta-analysis map of executive and cognitive control (see Materials and Methods). The activation location in the DP-task vs. D-task mostly overlapped with the meta-activation map in fronto-parietal regions (Fig. 5B). Region of interest (ROI) analysis further revealed significant activation within the meta-analysis maps during the 100% trials in DP task, relative to both D-task and DP-task with other Odds [Fig. 5C; DP-100% vs. D-task: t(24) = 2.3, P < 0.05; DP-100% vs. DP-others: t(24) = 3.6; P < 0.01]. However, relative to uncertain trials (10%), such a significant difference was not observed in uncertain trials (40% and 70%), in which reaction times were prolonged [t(24) = −0.99, P = 0.33]. Taken together, the brain activation in DP-task vs. D-task was most likely related to cognitive control.

Based on the involvement of cognitive control in DP-task, we made an *a posteriori* hypothesis that the paradoxical elongation of reaction time in 100% conditions in DP-task (Fig. 3A) is associated with the cost for strategy switching (Sakai, 2008). Indeed, the meta-analysis map of switching (Fig. 5D, left) was similar to that of cognitive/executive control (Fig. 5B), which is reasonable because switching is an executive function implicated in fronto-parietal mechanisms (Kim et al., 2012). ROI analysis for the meta-analysis maps also revealed significant activation during the 100% trials [Fig. 5E; DP-100% vs. D-task: t(24) = 2.9, P < 0.01; DP-100% vs. DP-others: t(24) = 3.2; P < 0.01]. Again, within this ROI, a significant difference was not observed between uncertain trials (40% and 70%) and relative to uncertain trials (10%) [t(24) = −0.99, P = 0.33]. Consistent results were obtained by using meta-analysis maps based on a more conservative testing procedure (Yarkoni et al., 2011)(Fig. S3).

If strategy switching explains the elongation of reaction time in the trials with 100% Odds in the DP-task, then strategy switching must have taken place within the DP-task, between 100% and other probability conditions. In line with this assumption, comparing 100% probability conditions and the other conditions in DP-task revealed brain activations in pIFC and SPL (Fig. 6A and Table S7). However, such a significant difference was not observed between uncertain trials (40% and 70%) and more certain trials (10% trials), whereas reaction times differed between these conditions.

In contrast, no significant brain activation was observed in the comparison between without delay conditions (“When” = “Now”) and the other conditions in DP-task (Fig. 6B). Furthermore, the meta-analysis map almost included the activation location during trials with 100% Odds relative to trials with other probabilities in DP-task (Fig. 5D, right). Thus, the meta-analysis provided support for the (*a posteriori*) hypothesis that participants switched strategy between 100% Odds and the other probability conditions in DP-task.

To ensure the functional interactions among these brain regions during strategic multifactor decision-making, we conducted psychophysiological analysis (PPI) (Friston et al., 1997) using six areas specifically recruited in DP-task relative to D-task (Fig. 6C). Compared with D-task, DP-task resulted in significantly positive PPI from OTC and SPC [t(24) = 3.08, P < 0.05, corrected], and PPI from OTC to pLPFC failed to survive the multiple comparisons, but were almost significant [t(24) = 2.11, P < 0.05, uncorrected]. These results suggest that the explicit presentation of “100%” probability to the participant activated OTC, and this in turn activated pIFC and SPC to execute strategy switching. Taken together, the results of behavior and functional imaging collectively suggest that the participants used multiple strategies to handle probabilistic uncertainty, but not delay, in a cognitively demanding DP-task.

## Discussion

In this study, we investigated the role of executive control mechanisms in a cognitively demanding gambling task in which participants were required to simultaneously consider two independent factors that modulate the value of the offers, namely probability and delay. Relative to control tasks, explicit processing of probability, but not delay, in the delayed gambling task (DP-task) biased the participants’ decisions. In the DP-task, reaction time was increased in 100% Odds conditions relative to the other Odds conditions. Neuroimaging analyses were then conducted to understand these behavioral results. Explicit processing of probability, but not delay, in DP-task resulted in brain-wide activations in the frontal, parietal, and occipitotemporal areas. The pattern of brain activations implicated the involvement of executive control, in particular strategy switching, which could explain the elongation of reaction time in the multifactor task. Relative to the cognitively less demanding single-factor control task (D-task and P-task), the DP-task upregulated functional connectivity between OCT to pIFC and OCT to SPC, consistent with the brain-wide state-reconfiguration associated with the strategy switching. Taken together, these results suggest that in cognitively demanding, multifactor decision-making tasks, the executive control mechanism implicated in fronto-parietal regions plays a key role in handling cognitively burdensome factors such as probability.

In DP-task as well as in the two control tasks, probability and delay both affected the participant’s accept rate. The change in accept rate most likely reflected a change in the subjective value of the offers. In both DP-task and D-task, the accept rate decreased significantly as the delay increased, consistent with delay discounting of subjective value in behavioral economics (Ainslie, 2005; Frederick et al., 2002b; Green et al., 1981; Green et al., 1994; Green et al., 1999; Kirby, 1997; Mischel et al., 1989; Rachlin et al., 1991). Similarly, in both the DP-task and the P-task, an increase in the probability to obtain the reward resulted in an increase of the accept rate, consistent with probability discounting of subjective values (Camerer, 1995; Green and Myerson, 2004; Kahneman and Tversky, 1979; Ostaszewski et al., 1998; Rachlin et al., 1991; Starmer, 2000; Tversky and Kahneman, 1992). Furthermore, the present results confirm previous findings that delay discounting and probability discounting simultaneously and jointly affect decision-making behavior when the two factors are present (Blackburn and El-Deredy, 2013; Vanderveldt et al., 2015). The consistency with previous studies also confirms that the participants performed value-based decision-making both in the DP-task and the two control tasks. Furthermore, we confirmed previous reports that the behavioral effect of probability is larger than that of delay (Blackburn and El-Deredy, 2013; Vanderveldt et al., 2015), corroborating the notion that humans do not simply add multiple factors when performing a complex value-based decision-making task. Larger and more widespread brain activations for processing probability relative to processing delay (Fig. S2A-B) are also in line with this notion. It remains unclear whether the larger activation related to probability is a “cause” or “consequence” of the asymmetric processing of probability and delay. Further research is needed to address this point.

A key difference between previous behavioral economics studies and the present study is that we used decoding of neuroimaging results to suggest potentially relevant mental function related to the effects of the presence of multiple factors in value-based decision-making. Previous studies in cognitive psychology have reported that humans use different cognitive strategies to solve cognitively demanding tasks vs. cognitively less demanding tasks (Payne et al., 1963). For example, when faced with cognitively demanding decision-making tasks, human participants do not treat all available information equally; instead, they use a heuristic to prioritize and take into account only a subset of information (Brandstätter et al., 2006). Such phenomena are not only observed in laboratory experiments but are also commonly seen in real-life situations (Galotti, 2007). However, although value-based decision-making in real-life often needs to take into account multiple variables (*e.g.*, investments), there has been little exploration of cognitive psychological aspects of the presence of multiple factors in value-based decision making. In the present study, by comparing multi- and single-factor decision-making tasks in a controlled manner, we found modulations of the participants’ decision-making behaviors that could not be explained by probability discounting or delay discounting (Festinger, 1954; Keeney, 2010; Rangel et al., 2008; Vanderveldt et al., 2015). Of note, the change in accept rate (Fig. 2) was similar or even larger compared with probability discounting or delay discounting. Thus, cognitive psychological effects may influence value-based decision-making in real-life as strongly as key factors discovered in behavioral economics.

It is possible to argue that the change in accept rate seen in the DP-task was due to the structure of the block design used in the present study (Fig. 1B). Accept rate is a function of subjective value that is not absolute but depends on a reference point that is determined based on the current expectation for gain (Abeler et al., 2011). In the D-task block, all the trials had 100% Odds. In contrast, in the DP-task block, only a subset of trials had 100% Odds. This design made the subset of trials in the DP-task with 100% Odds have larger subjective values than the rest of the trials in the same block. Thus, if the reference point is created within each block, the subjective value of trials with 100% Odds in a DP-task may be higher than the trials in a D-task, which in turn, causes a higher accept rate in DP-task. However, this explanation is unlikely because accept rates did not change between the DP-task with an immediate outcome (*i.e.*, “When” = “Now”) and the P-task, even though the accept rate should increase in the latter condition if this explanation were true. Thus, the task design is unlikely to affect the change in accept rate due to the number of factors observed in the present study.

The explicit processing of 100% Odds in the DP-task may engage the certainty effect (Tversky and Kahneman, 1986). Typical behavioral tasks related to the certainty effect are presented with two options: "winning $30 with 100% certainty" and "winning $45 with 80% certainty and winning nothing for 20% certainty." Participants typically chose the former option despite its lower expected value, suggesting that the certainty for reward acquisition increased the subjective value. However, because the two conditions in DP-task and D-task compared in the present study both had 100% Odds, the present observation was not an effect of unequal weighting to 100% Odds relative to the other probabilities. Moreover, the increase of reaction time despite the decrease in uncertainty in DP-tasks with 100% Odds cannot be explained by the certainty effect alone. Taken together, the certainty effect alone is unlikely to explain the modulation of decision-making behaviors in the DP-task.

Motivation to obtain the reward is an important factor determining value-based decision behaviors that was not explicitly manipulated in our study. Using a multifactor, value-based decision-making task in which effort and probability to obtain reward were simultaneously manipulated (effort expenditure for reward task, EEfRT; Treadway et al., 2009), Treadway and colleagues found that human subjects with weaker motivation for reward tend to choose an option with smaller reward and smaller required effort over an option with larger reward and larger required effort. Such (negative) correlation is present when the probability of winning is low, but is largely abolished when the probability is high. In contrast, in our study, we observed delay discounting regardless of the probability to obtain the reward (Fig. 2), suggesting that the cost associated with acceptance or rejection of offers was largely constant to the participants. Although we believe that this task condition was beneficial for isolating fMRI activity related to the processing of probability and delay *per se*, the downside was that we were unable to measure whether and how effort, and hence reward motivation, could have modulated the decision-making behaviors in our task. Rather than combining probability, delay, and effort in a single task, however, one can conduct a complex value-based decision task (e.g., DP-task) and EEfRT in separate sessions with the same participants to quantify reward motivation in individual participants (Giustiani et al., 2020). Such measurement is not only important for understanding the effect of individual personality on complex value-based decision-making but also for understanding how human participants with psychiatric disorders might respond to complex value-based decision tasks. Reward motivation strongly modulates decision-making behaviors of human participants with psychiatric disorders such as schizophrenia (Green et al., 2015; Huang et al., 2016). Thus, quantification of reward motivation would be indispensable to understand behaviors of participants with psychiatric disorders in complex value-based decision tasks.

Brain activity in the frontal and parietal regions related to the acceptance of gambles is consistent with previous results that brain activity increases in the frontal and parietal regions when expectations for rewards are high (Fig. S2C) (Hare et al., 2008; Rolls, 2000; Tom et al., 2007). The frontal and parietal activations related to the increase in probability can also be explained by the higher expectations for rewards. It should be noted that we did not detect significant activation in the orbitofrontal cortex (OFC), which is known as a brain region related to subjective value (Kable and Glimcher, 2007; Levy and Glimcher, 2012). This is most likely due to the fact that the present tasks were not designed to isolate the effect of subjective value (Jimura et al., 2011). Brain activity in the OTC increased due to the increase in the DP-task relative to the control tasks. Given that the region corresponds to early visual areas, the activation in the OTC most likely reflects additional visual processing due to the increased number of visual stimuli relative to the control tasks (Fig. 1A) (Mentis et al., 1997; Zeki, 1978).

Explicit processing of probability in the DP-task activated the pIFC, SPC, and OTC relative to the D-task. Activation in the OTC likely reflects an increase in visual information processing, as described above. The present meta-analyses revealed that activations in pIFC and SPC overlapped with brain regions previously implicated for strategy switching (Konishi et al., 2002; Sakai and Passingham, 2006). This pattern of brain activation suggested an interpretation that the participants switched their decision-making strategy to handle a subset of conditions (*i.e.* conditions with 100% Odds) in a cognitively demanding DP-task. In fact, pIFC and SPC were similarly activated in the 100% Odds conditions in DP-task relative to the other Odds conditions in DP-task. Consistently, the reaction time for 100% Odds tended to be longer than other Odds, likely reflecting the switching cost (Rogers and Monsell, 1995) associated with the shift in strategy between 100% Odds and the other Odds. It is intriguing that strategy switching did not seem to occur in the 10% Odds conditions, which also had a low probabilistic uncertainty. The differential processing of 100% and 10% Odds conditions may reflect the certainty effect (Tversky and Kahneman, 1986). It is of great interest which strategies the participants were switching between. Notably, a recent large-scale study reported that human participants frequently adopted flexible decisions that switched between subjective value-based decisions and expected utility-based decisions (Peterson et al., 2021). We speculate that the participants in the present study adopted similar flexible decisions during DP-task.

The decoding of functional maps showed strong weights for the terms related to cognitive/executive control including working memory and switching (Fig 5A). Working memory is a representative executive (cognitive) control function that refers to active maintenance and updating of goal-relevant information (D’Esposito and Postle 2015). Switching is also a representative executive control function, specifying shifting between one engagement to another. Thus, as executive control functions, switching and working memory partially share functional constructs, and indeed, it is well-known that cortical involvements in working memory and switching overlap in broad fronto-parietal regions (Dosenbach et al. 2006). Importantly, the decoder that we used to create the word cloud (Yarkoni et al. 2011) calculated association weights between activation maps and terms based on topographical similarity between the maps and meta-analysis maps related to the terms. Therefore, it is reasonable that both task (switch) and working memory shows strong weights.

Given the topographical similarity of brain regions involved in working memory and switching, theoretical accounts of these functions may help to specify cognitive processing in the current task. Working memory helps to guide decisions by encoding, maintaining, and integrating choice information. It has also been suggested that engagements of working memory are especially helpful for difficult decision-making in which choice information needs to be evaluated elaborately (Jimura et al. 2018). On the other hand, in the current study, we examined brain activity during decision-making, but comparisons were made based on logically equivalent situations (DP-100% and D-task trials) and easier situations (DP-100% vs. DP-others trials). Thus, working memory functioning is subtracted out in our comparisons for the DP-100% trials, and it is possible to interpret our activation results as not directly reflective of working memory.

Switching from one task to another requires cognitive processing to release a set of task rules and implement another set. Despite easier decision situations, prolonged reaction times in DP-100% are indicative of strategy switching. More importantly, our behavioral results suggest a switch of strategy when the DP-100% trials were performed. Taken together, this collective behavioral evidence and theoretical accounts suggest that switching may be more suitable to explain cognitive processing specifically involved in the DP-100% trials.

PPI analysis revealed increased functional connectivity from the OTC to pIFC and OTC to SPC in DP-task, with 100% Odds relative to D-task. Similarly, a small but significant negative change of functional connectivity was found from the pIFC to SPC. The location of pIFC activation in the present study is close to the site in pIFC implicated for feedback processing but not the site in pIFC implicated for response inhibition (Hirose et al., 2009). Thus, activations in the pIFC and SPC in the present study may represent different cognitive components involved in strategy switching (Crone et al., 2006; Derrfuss et al., 2005; Yeung et al., 2006).

It should be noted that the present conclusion related to strategy switching is based on the *a posteriori* hypothesis which was derived from the behavioral results that the reaction time was elongated in DP-100%. The *a posteriori* hypothesis was examined in order to provide a possible interpretation that could comprehensively explain the counterintuitive behavioral results together with brain activations. Although the test of a post-hoc hypothesis in general has a risk of reverse inference (Krajbich et al., 2015; Poldrack, 2006; Poldrack and Yarkoni, 2016), we took great care to avoid misinterpretations related to reverse inference. First, we carefully compared the present behavioral results with previous literature to make sure that the difference in reaction time between DP-task and the control task (from which we derived the *a posteriori* hypothesis) is unlikely to be an artifact of the task structure. Krajbitch and colleagues suggested that reaction time effects in behavioral economics experiments could artifactually arise simply from the fact that reaction time is expected to be longer when utilities for accept and reject are close to each other (Krajbich et al., 2015). In the present DP-task, we observed that reaction time was elongated in 100% Odds, despite the fact that the condition was designed to have the maximum difference of utility between accept and reject relative to other conditions. In contrast, reaction time in the control task showed short reaction time for 100% Odds relative to other conditions such as 40% and 70% Odds, consistent with the idea that a larger difference of the utility resulted in a faster reaction time (Fig. 3B). Second, we closely adhered to a recommendation that a reverse inference based on one data modality should be checked and verified by another data modality (Poldrack, 2006). According to this recommendation, the hypothesis of strategy switching which was derived from the behavioral data was tested on neuroimaging data. For the analysis of neuroimaging data, we employed a well-established and stringent large-scale, automated meta-analyses based on Neurosynth (Yarkoni et al., 2011). Furthermore, in addition to the stringent meta-analysis, we conducted PPI to confirm that brain activities were coordinated during DP-task, as described above. Taken together, we consider that the present conclusion is not affected by common pitfalls related to the use of reverse inference. Nevertheless, we are aware that these arguments do not fully eliminate the risk, and thus care needs to be taken to interpret the results related to the *a posteriori* hypothesis. Future studies using brain stimulation (Hill et al., 2017) or lesion studies (Noonan et al., 2017) are needed to further establish the involvement of strategy switching in complex value-based decision-making.

In the current task, participants made choices in hypothetical situations, which could affect affects participants’ seriousness and engagement with the task and which could be reflected in the current imaging results. However, when collecting data in a laboratory experiment where participants make choices based on the amount of monetary reward and years of delay to receive choice outcomes, hypothetical situations would be inevitable. Indeed, previous studies of value-based decision-making have used hypothetical situations when presenting rewards delayed by years (e.g., Rachlin et al., 1999; Green et al. 1994, 1999; Vanderveldt et al. 2015, Jimura et al. 2018).

To address this issue, some studies provided choice outcomes by randomly choosing from the trials, and participants received a payment after specified delay (e.g., Kable and Glimcher 2007). However, it is possible that such randomly chosen real outcomes might distort their engagements; participants would be engaged more in the trials where it was more realistic to receive money from the experimenter through the presented choice. We wished to circumvent this potential distortion in the current study.

Behaviorally, we obtained reasonable results of accept rates, consistent with a prior study (Tom et al. 2007), which assured that participants appropriately performed the task. Nonetheless, due to the nature of the hypothetical situations, the neural effects in our experiment might be weaker than those in the real choice.

## Acknowledgements

This study was supported by JSPS Kakenhi, 26350986, 26120711, 17H05957, 17K01989 to KJ, 17H00891 to KN; 20H00521 to MT; 19K16252, 20H05052 and 21H0516513 to TM; a grant from Brain/MINDS Beyond (AMED) to TM (grant number JP20dm0307031); a grant from JST-PRESTO to TM; a grant from Uehara Memorial Foundation to KJ; a grant from Takeda Science Foundation to KJ and MT; Keio Gijuku Academic Development Funds to KJ; and a grant from Keio Leading-edge Laboratory of Science and Technology to KJ. We thank Maoko Yamanaka for administrative assistance. We also thank Satoshi Hirano for technical assistance.

## Data and code availability statement

The originally collected data and developed scripts will be publicly available upon the publication of the current study via public repository.

## Supplementary Information

**Figure S1.**
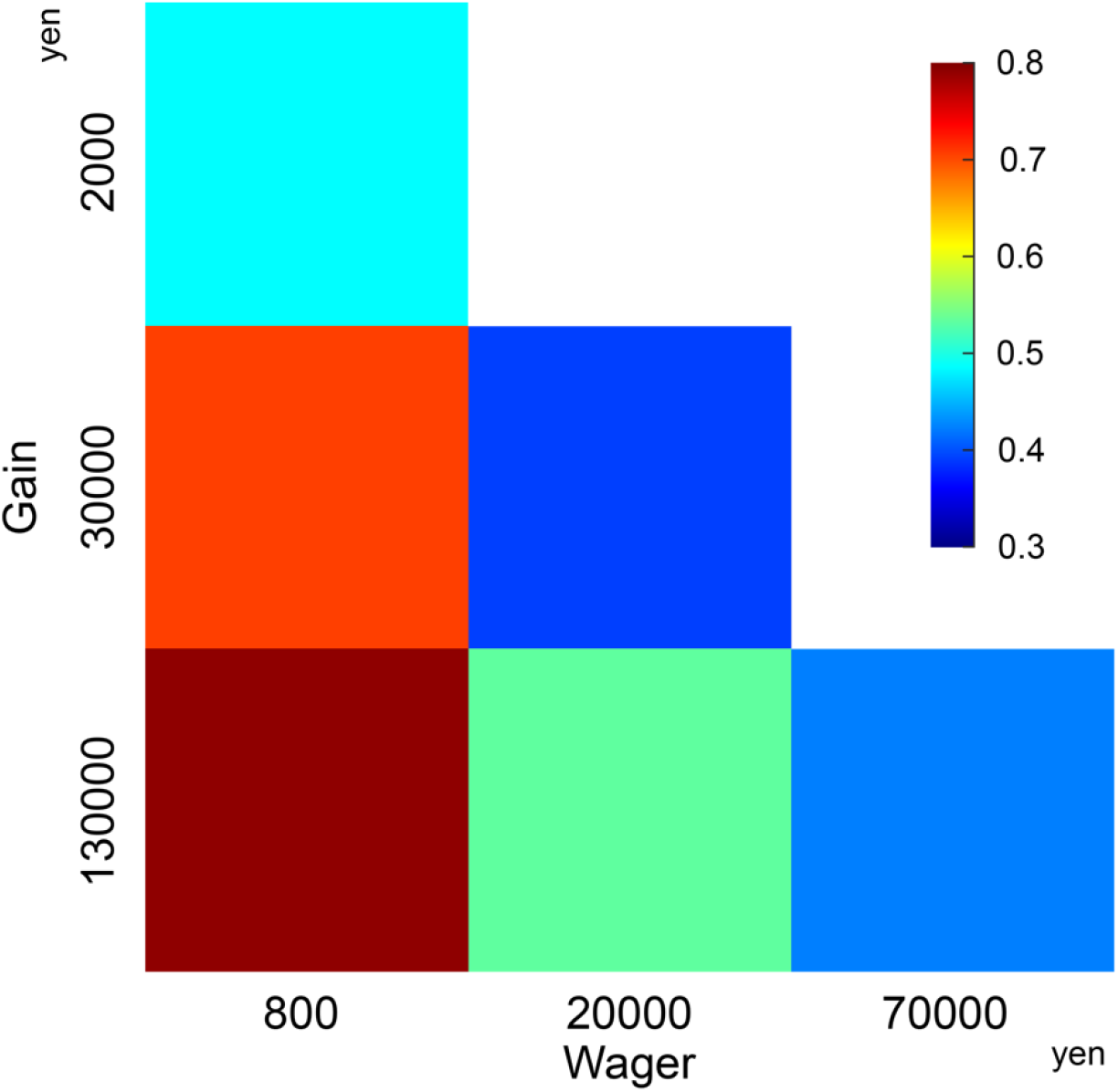
Accept rates in DP task as a function of Gain and Wager.

**Figure S2.**
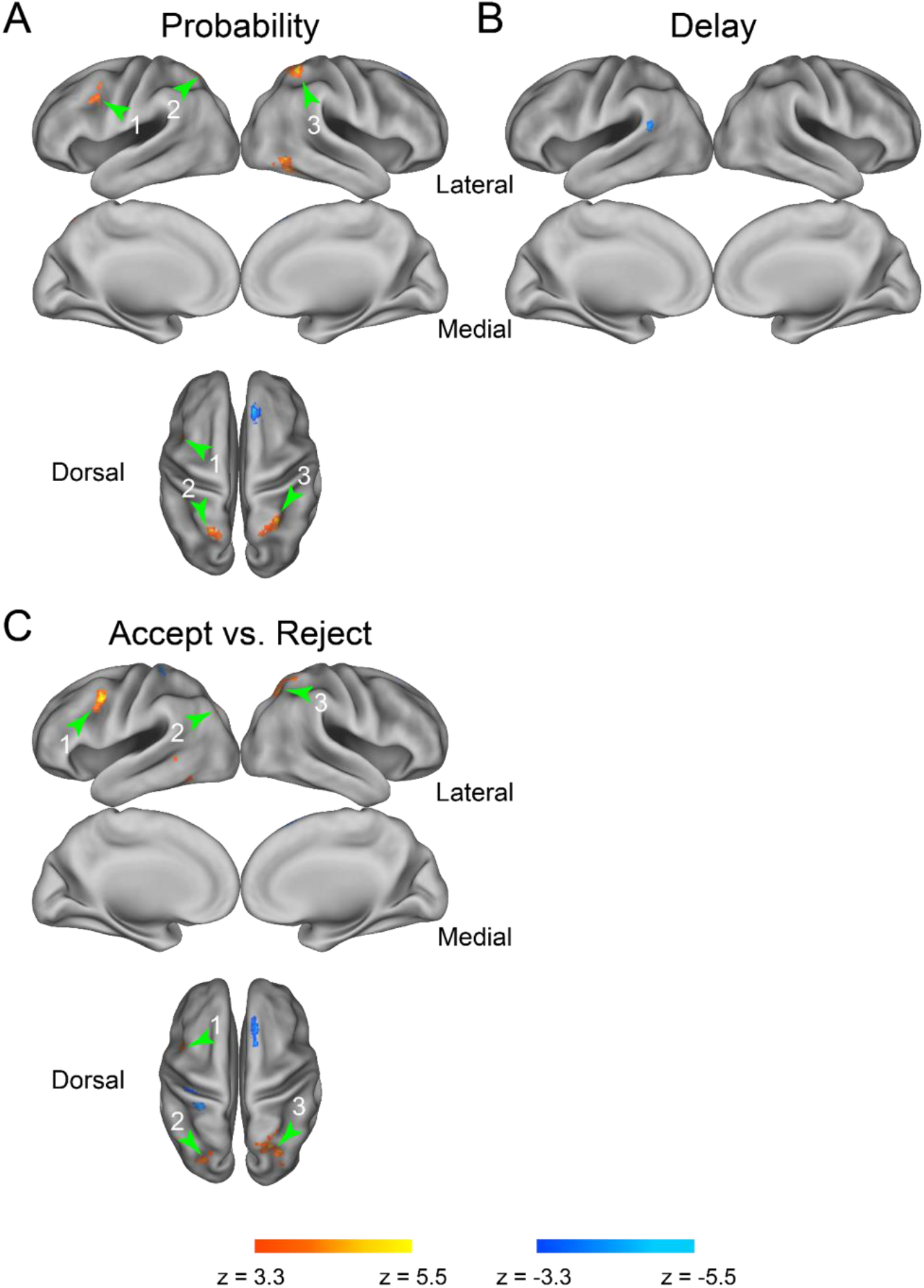
A/B) Statistical activation maps of brain regions showing activity modulation of probability (A) and delay (B) of the gamble. Maps were overlaid onto 3D surface of the brain. Hot and cool signal indicate greater activity in higher and lower probability/delay, respectively. C) Statistical activation maps of brain regions showing greater activity in trials in which the gamble was accepted (hot) and rejected (cool). 1: posterior lateral prefrontal cortex; 2/3: left/right superior parietal cortex.

**Figure S3.**
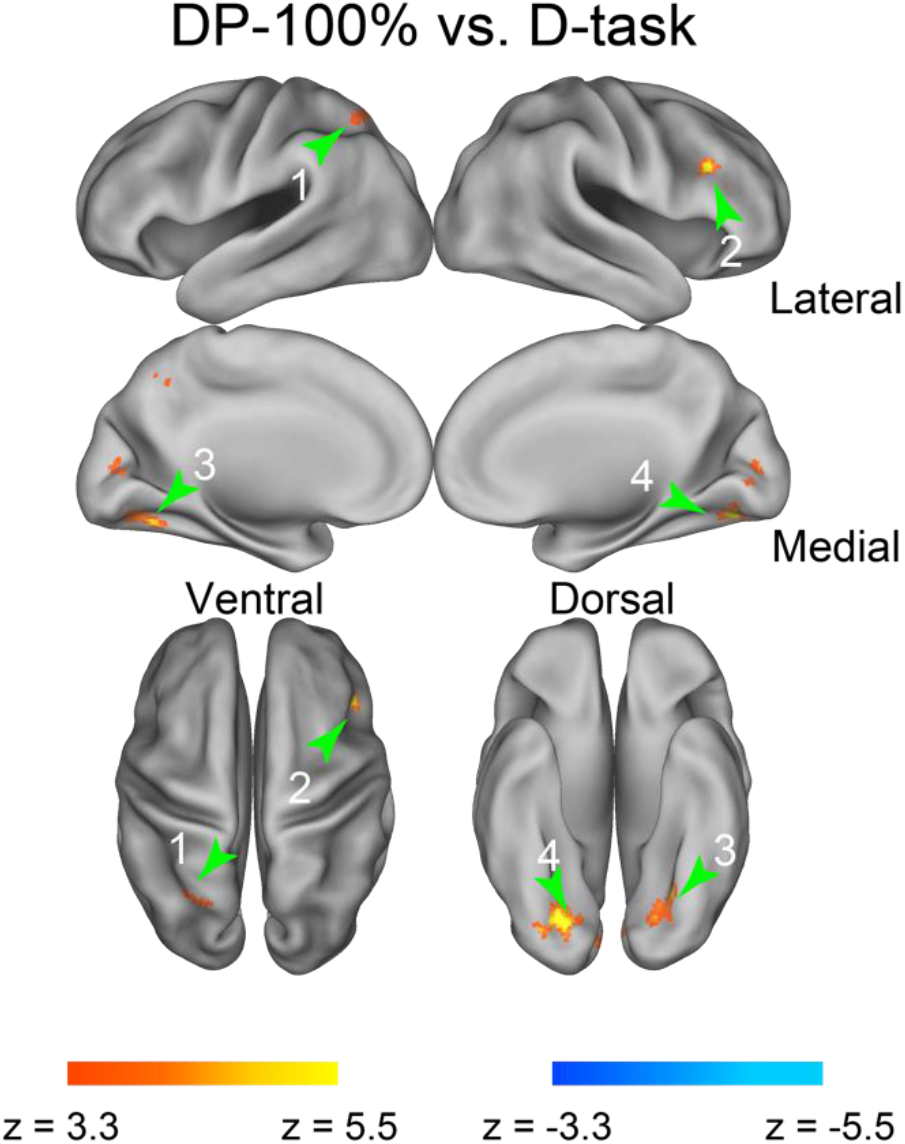
Brain activations related to processing of probability and delay in the multifactor task A) Activation related to probability processing in multifactor context. Reaction time of each trial coded as a parametric regressor in GLM analysis to minimize general difference in cognitive load. Statistical activation maps of brain regions showing greater activity in 100% Odds trials in DP-task (hot) and physically equivalent control trials (D-task) (cool). Formats are similar to those in Fig 4A.

**Figure S4.**
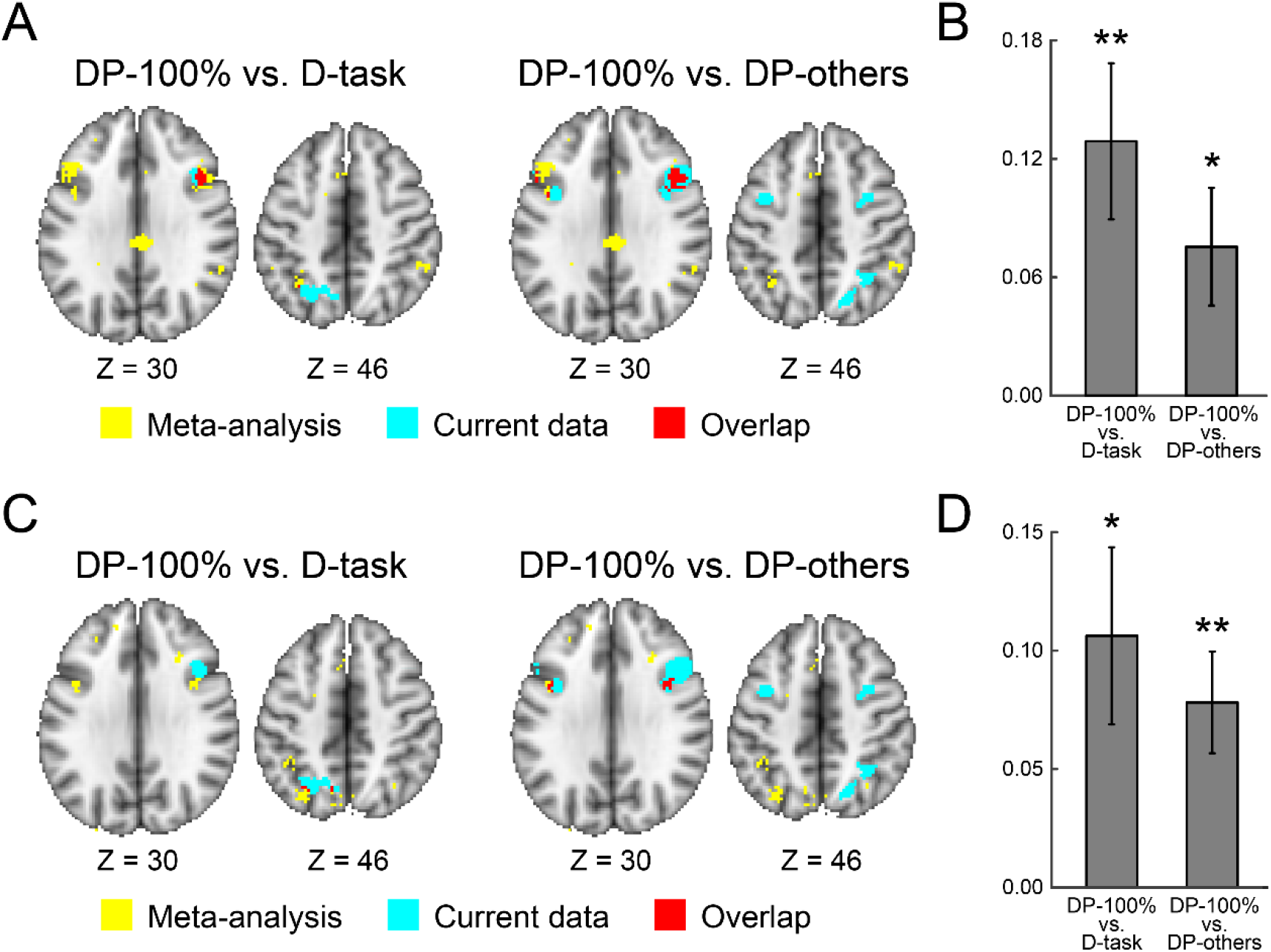
A) Maps of meta-analysis for cognitive/executive control with the association test are overlaid onto 2D transverse slices in yellow. Other formats are identical to those in Fig. 5B. B) Region of Interest (ROI) analysis. ROI was defined based on the meta-analysis maps in panel A, and other formats were identical to those in Fig. 5C. DP-100% vs. D-task: t(24) = 3.2, P < 0.01; DP-100% vs. DP-others: t(24) = 2.5, P < 0.05. C) Maps of the meta-analysis for switching with the association test are overlaid onto 2D transverse slice in yellow. Other formats are identical to those in panel A. D) ROI analysis. ROIs were defined based on the meta-analysis maps in panel C, and other formats were identical to those in panel B. DP-100% vs. D-task: t(24) = 2.3 P < 0.05; DP-100% vs. DP-others: t(24) = 3.6, P < 0.01.

**Table S1.** Brain regions showing significant parametrical effect with probability. Coordinates are listed in MNI space. Positive and negative t-values indicate increase in high and low probability, respectively. BA indicate Brodmann areas and is approximate. Cluster size is in voxels.

| Area | Coordinate |  |  | t-value | BA | Cluster |  |
| --- | --- | --- | --- | --- | --- | --- | --- |
|  | x | y | z |  |  | ID | Size |
| Frontal | -38 | 2 | 42 | 4.63 | 9/6 | 1 | 136 |
|  | 10 | 32 | 56 | -6.15 | 8 | 2 | 259 |
|  | 12 | 20 | 64 | -4.77 | 8 | 2 | 259 |
| Parietal | 24 | -56 | 54 | 5.34 | 7 | 3 | 296 |
|  | 16 | -64 | 58 | 4.22 | 7 | 3 | 296 |
|  | -20 | -64 | 54 | 4.90 | 7 | 4 | 131 |
| Occipitotemporal | 46 | -60 | -8 | 5.86 | 37 | 5 | 212 |
|  | 44 | -72 | -10 | 3.78 | 37 | 5 | 212 |

**Table S2.** Brain regions showing significant parametrical effect with delay duration. Positive and negative t-values indicate increase in long and short delay, respectively. Formats are similar to those in Table S1.

| Area | Coordinate |  |  | t-value | BA | Cluster |  |
| --- | --- | --- | --- | --- | --- | --- | --- |
|  | x | y | z |  |  | ID | Size |
| Temporal | -58 | -52 | 16 | -5.19 | 39 | 1 | 132 |

**Table S3.** Brain regions showing significant signal increase and decrease in the contrast accept versus reject decisions. Positive and negative t-values indicate increase in accept and reject decisions, respectively. Formats are similar to those in Table S1.

| Area | Coordinate |  |  | t-value | BA | Cluster |  |
| --- | --- | --- | --- | --- | --- | --- | --- |
|  | x | y | z |  |  | ID | Size |
| Frontal | -42 | 0 | 40 | 5.64 | 6/9 | 1 | 142 |
|  | 12 | 10 | 62 | -5.32 | 6 | 2 | 199 |
|  | 14 | 22 | 62 | -5.06 | 6/8 | 2 | 199 |
| Parietal | 22 | -62 | 48 | 5.49 | 7 | 3 | 426 |
|  | 26 | -62 | 60 | 5.01 | 7 | 3 | 426 |
|  | 26 | -62 | 60 | 4.96 | 7 | 3 | 426 |
|  | 30 | -76 | 30 | 3.88 | 39 | 3 | 426 |
|  | -28 | -74 | 28 | 4.59 | 39 | 4 | 143 |
|  | -22 | -66 | 38 | 4.08 | 39 | 4 | 143 |
|  | -30 | -62 | 26 | 4.01 | 39 | 4 | 143 |
|  | -40 | -28 | 48 | -4.26 | 5 | 5 | 216 |
|  | -28 | -38 | 60 | -5.91 | 5 | 5 | 216 |
| Occipitotemporal | -46 | -56 | -14 | 4.65 | 37 | 6 | 125 |
|  | -60 | -50 | 0 | 4.06 | 36/37 | 6 | 125 |

**Table S4.** Brain regions showing significant signal increase and decrease between trials with 100% Odds in DP-task and D-task trials. Positive and negative t-values indicate increase in trials with 100% Odds and D-task trials, respectively. Formats are similar to those in Table S1.

| Area | Coordinate |  |  | t-value | BA | Cluster |  |
| --- | --- | --- | --- | --- | --- | --- | --- |
|  | x | y | z |  |  | ID | Size |
| Frontal | 42 | 16 | 30 | 5.90 | 44 | 1 | 139 |
|  | 54 | 24 | 38 | 3.85 | 8/6 | 1 | 139 |
| Parietal | -14 | -60 | 46 | 5.29 | 7 | 2 | 231 |
|  | -26 | -62 | 44 | 4.66 | 7 | 2 | 231 |
|  | -6 | -68 | 42 | 3.77 | 7 | 2 | 231 |
| Occipitotemporal | -20 | -70 | -12 | 5.67 | 18 | 3 | 245 |
|  | -32 | -72 | -10 | 5.06 | 18 | 3 | 245 |
|  | 4 | -84 | 4 | 4.98 | 17 | 4 | 410 |
|  | 14 | -74 | 10 | 4.65 | 17 | 4 | 410 |
|  | -12 | -80 | 8 | 4.46 | 17 | 4 | 410 |
|  | -4 | -66 | 6 | 4.11 | 17 | 4 | 410 |
|  | 22 | -74 | -10 | 4.93 | 18 | 5 | 129 |
|  | 26 | -82 | -18 | 3.77 | 18/19 | 5 | 129 |

**Table S5.** Brain regions showing significant signal increase and decrease between trials with 100% Odds in DP-task and D-task trials. Reaction time of each trial coded as a parametric regressor in GLM analysis to minimize general difference in cognitive load. Positive and negative t-values indicate increase in trials with 100% Odds and D-task trials, respectively. Formats are similar to those in Table S1.

| Area | Coordinate |  |  | t-value | BA | Cluster |  |
| --- | --- | --- | --- | --- | --- | --- | --- |
|  | x | y | z |  |  | ID | Size |
| Frontal | 42 | 16 | 30 | 5.84 | 44 | 1 | 130 |
| Parietal | -14 | -60 | 46 | 4.92 | 7 | 2 | 151 |
|  | -28 | -60 | 46 | 4.38 | 7 | 2 | 151 |
| Occipitotemporal | 22 | -74 | -10 | 7.05 | 18 | 3 | 265 |
|  | -20 | -70 | -12 | 5.29 | 18 | 4 | 251 |
|  | -32 | -72 | -12 | 5.12 | 18 | 4 | 251 |
|  | 4 | -84 | 4 | 4.82 | 17 | 5 | 157 |
|  | -12 | -82 | 8 | 4.32 | 17 | 5 | 157 |

**Table S6.**
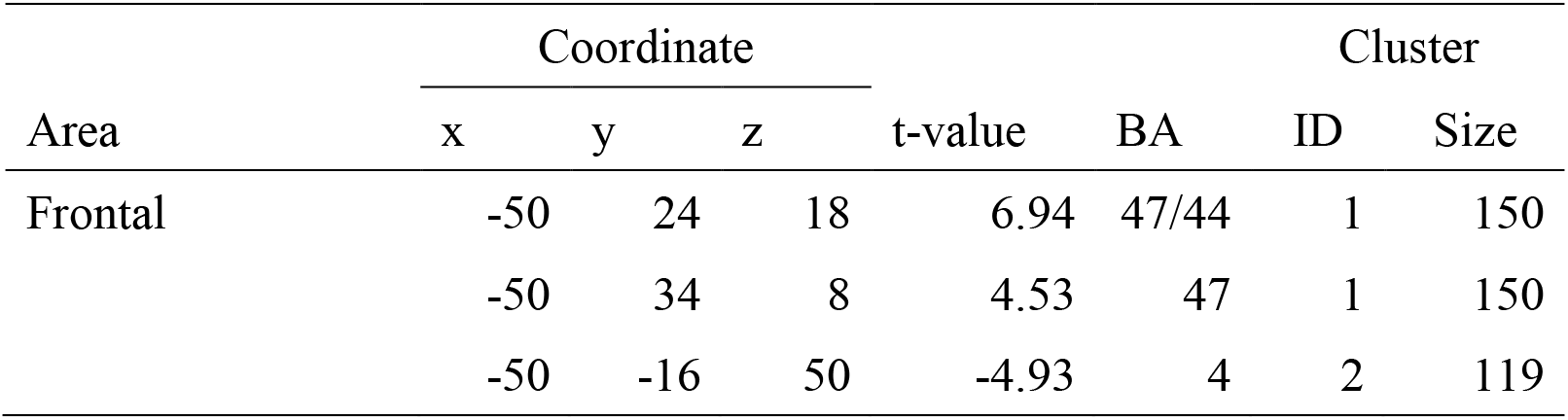
Brain regions showing significant signal increase and decrease between Now trials in DP task and P-task trials. Positive and negative t-values indicate increase in Now trials and P-task trials, respectively. Formats are similar to those in Table S1.

| Area | Coordinate |  |  | t-value | BA | Cluster |  |
| --- | --- | --- | --- | --- | --- | --- | --- |
|  | x | y | z |  |  | ID | Size |
| Frontal | -50 | 24 | 18 | 6.94 | 47/44 | 1 | 150 |
|  | -50 | 34 | 8 | 4.53 | 47 | 1 | 150 |
|  | -50 | -16 | 50 | -4.93 | 4 | 2 | 119 |

**Table S7.** Brain regions showing significant signal increase and decrease between trials with 100% Odds and trials with the other probabilities in DP-task. Positive and negative t-values indicate increase in trials with 100% Odds and trials with the other probabilities, respectively. Formats are similar to those in Table S1.

| Area | Coordinate |  |  | t-value | BA | Cluster |  |
| --- | --- | --- | --- | --- | --- | --- | --- |
|  | x | y | z |  |  | ID | Size |
| Frontal | -38 | 4 | 34 | 6.49 | 8 | 1 | 300 |
|  | -50 | 16 | 32 | 3.80 | 8/9 | 1 | 300 |
|  | 52 | 14 | 28 | 5.75 | 44 | 2 | 664 |
|  | 38 | 22 | 22 | 5.73 | 46 | 2 | 664 |
|  | 32 | 2 | 46 | 5.39 | 4/6 | 2 | 664 |
|  | 38 | 4 | 32 | 4.36 | 4/6 | 2 | 664 |
|  | 58 | 26 | 24 | 4.22 | 44 | 2 | 664 |
|  | 48 | 10 | 40 | 4.08 | 6/4 | 2 | 664 |
| Parietal | 32 | -58 | 62 | 5.13 | 7 | 3 | 535 |
|  | 34 | -52 | 46 | 4.97 | 7 | 3 | 535 |
|  | 18 | -66 | 56 | 4.41 | 7 | 3 | 535 |
|  | 40 | -48 | 62 | 4.12 | 7 | 3 | 535 |

